# Modelling idiopathic intracranial hypertension in rats: contributions of high fat diet and testosterone to intracranial pressure and cerebrospinal fluid production

**DOI:** 10.1101/2023.01.31.526446

**Authors:** Jonathan H. Wardman, Mette Nyholm Jensen, Søren Norge Andreassen, Bjarne Styrishave, Jens E. Wilhjelm, Alexandra J. Sinclair, Nanna MacAulay

## Abstract

**Background:** Idiopathic intracranial hypertension (IIH) is a condition characterized by increased intracranial pressure (ICP), impaired vision, and headache. Most cases of IIH occur in obese women of childbearing age, though age, BMI, and female sex do not encompass all aspects of IIH pathophysiology. Systemic metabolic dysregulation has been identified in IIH with a profile of androgen excess. However, the mechanistic coupling between obesity/hormonal perturbations and cerebrospinal fluid dynamics remains unresolved.

**Methods:** Female Wistar rats were either fed a high fat diet (HFD) or exposed to adjuvant testosterone treatment to recapitulate IIH causal drivers. Cerebrospinal fluid (CSF) and blood testosterone levels were determined with mass spectrometry, ICP and CSF dynamics with *in vivo* experimentation, and the choroid plexus function revealed with transcriptomics and ex *vivo* isotope-based flux assays.

**Results:** HFD-fed rats presented with increased ICP, which was not accompanied by altered CSF dynamics or modified choroid plexus function. Chronic adjuvant testosterone treatment of lean rats caused elevated CSF secretion rate, in association with increased activity of the choroid plexus Na^+^,K^+^,2Cl^-^ cotransporter, NKCC1.

**Conclusions:** HFD-induced ICP elevation in experimental rats did not originate from an increased rate of CSF secretion. Such modulation of CSF dynamics only came about with adjuvant testosterone treatment, mimicking the androgen excess observed in female IIH patients. Obesity-induced androgen dysregulation may thus play a crucial role in the disease mechanism of IIH.

## INTRODUCTION

Idiopathic intracranial hypertension (IIH) is characterized by an unexplained increase in intracranial pressure (ICP) resulting in life-altering symptoms. These include severe, persistent headache and optic nerve swelling (papilloedema) that can lead to permanent visual loss and cognitive impairment [1–6]. Obese women of childbearing age make up approximately 90% of all IIH cases, though IIH can rarely afflict males, diverse ages, and body-types [5–8]. Increased global rates of obesity have led to increased IIH incidence (by greater than 350% in the last decade) [7, 9–11]. Visual impairments have improved due to increased awareness and earlier intervention [11–13], but severe headache and cognitive dysfunction continue to threaten those suffering from IIH, which can lead to long-term disability [3, 14, 15]. Pharmacological interventions for IIH are currently limited to unlicensed used of diuretic drugs such as acetazolamide and topiramate [6], which are comprised by side-effects and low adherence [2]. Severe IIH, with rapidly progressive visual loss, remains treatable only via surgical procedures such as ventriculo-peritoneal shunting, which is associated with marked complications [2]. Weight loss via dietary restriction or bariatric surgery has proven effective for reducing ICP and inducing remission in IIH [16, 17]. This apparent link between weight loss and symptom alleviation indicates a connection, currently unresolved, between obesity and the etiology of IIH [17].

The elevated ICP in IIH has been attributed to multiple factors, including aspects of CSF dynamics such as elevated brain fluid content [18–20] due to inefficient CSF clearance [21–24], increased CSF production rate [21, 25], or venous sinus stenosis [26, 27], as well as reduced compliance of brain tissue and vasculature due to fibrogenesis and deterioration of the basement membrane, among others [28]. Most of these suggested etiologies have, however, been contested [29–31]. Despite a clear link between female sex, obesity, and age and the symptoms of IIH, IIH only affects up to 15.2/100,000 patients fitting these criteria [7], and IIH is not endemic in the obese female population, suggesting the presence of other factors leading to emergent IIH. A range of factors of putative importance for disease etiology has been detected in IIH patients [32–34]. Most notably, analysis of IIH steroid hormone profiles revealed a unique phenotype of serum and CSF androgen excess, distinct from that driven by obesity alone [33]. The interplay between adiposity and androgen excess, both features of systemic metabolic dysregulation, and ICP elevation is not understood.

Obesity and androgen profiles are not readily manipulated in humans in order to determine their individual impact on ICP, brain fluid dynamics, and choroid plexus function. We therefore aimed to resolve the role of obesity and androgen excess on CSF dynamics in an animal model. To this end, we employed a high fat diet (HFD)-fed rat model mimicking aspects of IIH [35, 36] and a complementary rat model of elevated testosterone for determination of the ensuing brain fluid dynamics to elucidate their potential contribution to the ICP elevation that defines IIH.

## METHODS

### Experimental Animals

Animal handling and experiments were performed according to European guidelines and complied with all ethical regulations. It was approved by the Danish Animal Experiments Inspectorate with permission no. 2018-15-0201-01515. Female Wistar rats (Janvier Labs), aged 6 weeks, were divided into two groups of equal starting weight (HFD = 171 ± 10 g, n = 17, control = 173 ± 9, n = 14, p = 0.98) and were fed either a high fat diet (HFD, 60% Kcal from fat, Research Diets: D12492i) or a nutrient source-matched control diet (10% Kcal from fat, Research Diets: D12450ji) for 21 weeks. Diet and water were provided *ad libitum* and the rat weight recorded on a weekly basis. Experimental animals were chosen based upon weight, selecting control diet-fed rats weighing below a 305 g threshold and HFD-fed rats above a 335 g threshold, providing an 18% average weight difference between groups, similarly to weight differences and thresholds obtained in a previous study [35]. For testosterone experiments, female Wistar rats (Janvier Labs) were used at nine weeks of age, with either twice-weekly subcutaneous injections starting four weeks prior with 1 mg testosterone propionate in 100 μl sesame oil (86541, Sigma-Aldrich) [37] or 100 μl sesame oil (S3547, Sigma-Aldrich, vehicle control). Testosterone administration was calculated to obtain supraphysiological testosterone levels concomitant with those observed in female IIH patients [33, 37].

### Solutions

The majority of the experiments were conducted in CO_2_/HCO_3_^-^-buffered artificial cerebrospinal fluid (HCO_3_^-^-aCSF; (in mM) 120 NaCl, 2.5 KCl, 2.5 CaCl_2_, 1.3 MgSO_4_, 1 NaH_2_PO_4_, 10 glucose, 25 NaHCO_3_, equilibrated with 95% O_2_/5% CO_2_ to obtain a pH of 7.4). In experiments where the solution could not be equilibrated with 95% O_2_/5% CO_2_, as in isotope influx and efflux experiments, aCSF was instead buffered by HEPES (HEPES-aCSF; (in mM) 120 NaCl, 2.5 KCl, 2.5 CaCl_2_, 1.3 MgSO_4_, 1 NaH_2_PO_4_, 10 glucose, 17 Na-HEPES, adjusted to pH 7.4 with NaOH).

### Anesthesia and ventilation

Anesthesia was implemented via intraperitoneal (i.p.) injection of 6 mg/ml xylazine + 60 mg/ml ketamine (ScanVet) in sterile water (0.17 ml/100 g bodyweight, pre-heated to 37° C). Animals were re-administered half ketamine dose as required to sustain anesthesia. One rat was excluded because it was unresponsive to initial anesthesia administration. The body temperature was maintained at 37 °C by a homeothermic monitoring system with heat pad (Harvard Apparatus). Mechanical ventilation was employed for all anesthetic protocols lasting more than 30 min, to ensure stable respiratory partial pressure of carbon dioxide (pCO_2_) and oxygen (pO_2_) and arterial oxygen saturation and thus stable plasma pH and electrolyte content. Surgical tracheotomy was carried out for mechanical ventilation, which was controlled by the VentElite system (Harvard Apparatus) by 0.9 l/min humidified air mixed with 0.1 l/min oxygen adjusted with approximately 2.6 ml per breath, 80 breath/min, a Positive End-Expiratory Pressure (PEEP) at 2 cm, and 10 % sight for a ~350 g rat. Ventilation settings were optimized for each animal using a capnograph (Type 340, Harvard Apparatus) and a pulse oximeter (MouseOx^®^ Plus, Starr Life Sciences) after system calibration with respiratory pCO_2_ (4.5 – 5.0 kPa) and pO_2_ (13.3 - 17.3 kPa) and arterial oxygen saturation (98.8 - 99.4 %) (ABL90, Radiometer).

### ICP measurements

Anesthetized and ventilated rats, placed in a stereotactic frame, had the skull exposed, and a 3.6⍰mm diameter cranial window drilled with care not to damage the dura. The epidural probe (PlasticsOne, C313G) was secured with dental resin cement (Panavia SA Cement, Kuraray Noritake Dental Inc.) above the dura and the ICP probe was filled with HEPES-aCSF before connection to a pressure transducer APT300 connected to a transducer amplifier module TAM-A (Hugo Sachs Elektronik) followed by a multifunction data acquisition module DT9836-12-2-BNC (Data Translation). To ensure the presence of a continuous fluid column between the dura and the epidural probe, approximately 5⍰μl HEPES-aCSF was injected through the epidural probe. The ICP signal was recorded at a 1⍰kHz sampling rate using BDAS Basic Data Acquisition Software (Hugo Sachs Elektronik). Jugular compression was applied to confirm proper ICP recording.

### ICP Waveform analysis

The raw ICP data stored as semicolon separated values in normal text files by the BDAS software were read into MATLAB (MathWorks), and possible spikes and non-physiological data points removed. This was followed by low-pass filtering (anti-aliasing) and downsampling to 100 Hz. These signals were of length about 300 s to 900 s and contained approximately 1.3 Hz frequency components from the artificial ventilation, the heart rate as well as other minor interferences. Since the heart rate was in the region 2.5 to 5 Hz, the signal was spectrally bandpass filtered between 1.6 Hz and 5.5 Hz with a Tukey window. The instantaneous peak-to-peak amplitude was finally extracted from the filtered time signals and employed to obtain the mean wave amplitude.

### Brain water quantification

For brain water determination, rats were decapitated under anesthesia. The brain was rapidly dissected into a pre-weighed porcelain evaporating beaker (Witeg) and weighed within 1 min after brain isolation. The brain was then homogenized with a spatula (to increase surface area) and left at 100°C for approximately 90 hours to dry. After drying, the brain was weighed again and the difference in the two measurements corresponded to the brain water (in g) and was employed to obtain the water percentage.

### Ventriculo-cisternal perfusion

Rats were anesthetized, ventilated, and an infusion cannula (Brain infusion kit 2, Alzet) was stereotactically placed in the right lateral ventricle. A 0.5⍰mm (diameter) burr hole was drilled (1.3⍰mm posterior, 1.8⍰mm lateral to bregma), and a 4⍰mm (length) brain infusion cannula (Brain infusion kit2, Alzet) was glued in place on the cranium with the cannula placed into the lateral ventricle, through which a pre-heated (37°C, SF-28, Warner Instruments) HCO_3_^-^-aCSF containing 1⍰mg/ml TRITC-dextran (tetramethylrhodamine isothiocyanate-dextran, MW⍰=⍰I50,000; T1287, Sigma) was infused at 9⍰μl/min. CSF was sampled from cisterna magna at 5⍰min intervals with a glass capillary (30–0067, Harvard Apparatus pulled by a Brown Micropipette puller, Model P-97, Sutter Instruments) placed at a 5° angle (7.5⍰mm distal to the occipital bone and 1.5⍰mm lateral to the muscle-midline). The cisterna magna puncture and continuous fluid sampling prevents elevation of ICP during the procedure. The fluorescent content of CSF outflow was measured in triplicate on a microplate photometer (545⍰nm, SyneryTM Neo2 Multi-mode Microplate Reader; BioTek Instruments), and the CSF secretion rate was calculated from the equation [38]:

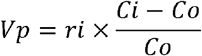

where *V_p_* = CSF secretion rate (μl/min), *r_i_* = infusion rate (μl/min), *C_i_* = fluorescence of inflow solution, *C_o_* = fluorescence of outflow solution.

### Live imaging of CSF movement

Rats were anesthetized, placed in a stereotactic frame and through a burr hole in the lateral ventricle (same coordinates as for ICP and ventriculo-cisternal perfusion) a Hamilton syringe (RN 0.40, G27, a20, Agntho’s) was placed (4 mm deep) with 15⍰μl HCO_3_^-^-aCSF with 10⍰μM carboxylate dye (MW=l,09l, IRDye 800 CW, P/N 929-08972, LI-COR Biosciences). For the acute testosterone treatment, an additional Hamilton syringe containing 15 μl HCO_3_^-^-aCSF with 2.5⍰μM testosterone (or 0.0025% DMSO as vehicle control) was injected into the ventricle five minutes prior the dye injection. The rat was swiftly placed in a Pearl Trilogy Small Animal Imaging System (LI-COR Biosciences) and within 1 min after ventricular dye injection, images were obtained at 30 s intervals (800⍰nm channel, 85⍰μm resolution, for 5 min). A white field image was acquired at the termination of each experiment, after which the rat was sacrificed. The isolated brain was then bisected to expose the ventricles to record a final micrograph ensuring proper targeting of the ventricular compartment. Images were analyzed in a blinded fashion using LI-COR Image Studio 5.2 (LI-COR Biosciences) and data presented as fluorescence intensity in a region of interest placed in line with lambda, normalized to the signal obtained in the first image.

### Androgen quantification in CSF and plasma

Rats were anesthetized, placed in a stereotactic frame and CSF was extracted through a cisterna magna puncture and immediately centrifuged to pellet cell debris (2000 × *g*, 10 min, 4°C) prior to storage of the ~ 100 μl supernatant at −80°C in sealed microcentrifuge tubes. Immediately after euthanization of each rat, a blood sample was collected into heparin-coated Eppendorf tubes and centrifuged (7500 × *g*, 5 min, 4°C). The ~ 400 μl plasma was transferred to new vials and stored at −80°C until analysis. The steroid extraction was performed as earlier described for plasma [39] with a further modification of the CSF analysis to encompass a different size of solid-phase extraction columns (100 mg Bond elute C_18_ solid-phase extraction cartridges; 1 ml; Agilent) and therefore corresponding volume changes for conditioning (1 ml MeOH followed by 2×1 ml dH_2_O), washing (2×1 ml dH_2_O followed by 1 ml H_2_O:MeOH (3:1)) and elution (1 ml H_2_O:MeOH (1:4)). The liquid chromatography online clean-up, chromatographic separation of androgens and mass spectrometry data analysis were done as earlier described [39].

### RNA sequencing

Choroid plexus (lateral and 4^th^) was isolated and stored in RNAlater^®^ (Sigma) at −80 °C prior to RNA extraction and library preparation with NEB Next^®^ Ultra^™^ RNA Library Prep Kit (NEB) by Novogene. RNA sequencing (paired-end 150 bp, with 12 Gb output) was performed on an Illumina NovaSeq 6000 (Illumina). All program parameter settings for library building and mapping, together with all scripts for the gene annotation and analysis are available at https://github.com/Sorennorge/-MacAulayLab-RNAseq3-Wistar. Raw data are available at the National Center for Biotechnology Information (NCBI) Gene Expression Omnibus (GEO) database (GSE223582). The sequencing data of 150 base paired-end reads were mapped to reference genome (Rattus norvegicus Rnor_6.0 v.104) using Spliced Transcripts Alignment to a Reference (STAR) RNA-seq aligner (v. 2.7.9a) [40]. The mapped alignment by STAR was both converted to raw counts from STAR GeneCounts and normalized to TPM with RSEM (RNA-Seq by Expectation Maximization v. 1.3.3) {Li, 2011 #199}[ref] The raw counts from STAR GeneCount were used for differential expression analysis using R library and program DEseq2 [41]. Differentially expressed genes were determined based on standard procedure of DEseq2 analysis with false discovery rate (FDR, Benjamini–Hochberg method) [42] of less than 0.05 [43]. The Volcano plot was created using R library ggplot2 [44], the subplot for the pie chart was created using python library matplotlib, and the subplot for Heatmap was generated using R library pheatmap [45]. The Gene Ontology (GO) enrichment analysis was generated utilizing the Panther database [46] with the gene symbols from the differentially expressed genes from DEseq2 to classify the protein class of each gene and the pie chart of the GO enrichment analysis created using python library matplotlib. The network analysis was generated from differentially expressed genes from DEseq2 using the gene symbols as protein database query from String-database (https://string-db.org/) [47] and only including connections with a string confidence score above 0.7 as a plugin for Cytoscape (v. 3.9.1) [48].

### ^86^Rb^+^ influx and efflux

Lateral choroid plexus was isolated from control and testosterone-treated rat brains and placed in 37°C HEPES-aCSF for a 5-10-min recovery period followed by 2 min (influx) or 8 min (efflux, with inclusion of bumetanide (20 μM) or vehicle) of incubation in an isotope solution containing rubidium (^86^Rb^+^) (1 μCi/ml, 022-105721-00321-0001, POLATOM) and ^3^H-mannitol (4 μCi/ml, NET101, Perkin Elmer). ^86^Rb^+^ acts as a K^+^ congener, and can be transported by the Na^+^, K^+^, 2Cl^-^ cotransporter, NKCC1, and the Na^+^/K^+^-ATPase, amongst others, in place of K^+^, whereas ^3^H-mannitol remains outside, serving as an extracellular marker [49]. The ^86^Rb^+^ transport rate is independent of whether the aCSF is buffered by HCO_3_^-^ or HEPES [50] and the experiments thus conducted in HEPES-buffered aCSF to avoid the required equilibration of the HCO_3_^-^-buffered aCSF, which may induce spraying of isotopes. For efflux assays, choroid plexuses were, in a paired fashion, randomly assigned to either control or bumetanide group (20 μM, Sigma, B3023, stock solution prepared in DMSO (Sigma D8418) prior to dilution in HEPES-aCSF to a final DMSO concentration of 0.1% as also employed as vehicle). For influx assays, choroid plexuses were, in a paired fashion, randomly assigned to either control or ouabain group (2 mM, Sigma, O3125, dissolved directly into HEPES-aCSF on the day of experiment). The acute exposure to testosterone consisted of a one hour preincubation of the choroid plexus in 37°C HEPES-aCSF containing 100 nM testosterone (dissolved in ethanol prior to dilution in HEPES-aCSF to a final vehicle concentration of 0.04%ethanol, which was employed in the control group) prior to initiation of the flux assay. In influx assays, choroid plexus was subsequently rinsed in ice-cold isotope-free HEPES-aCSF containing 2 mM ouabain, 20 μM bumetanide, and 1 mM BaCl_2_ (to prevent efflux of intracellular [^86^Rb^+^] during the washing procedure), followed by transfer to scintillation vials containing 100 μl Solvable (6NE9100, Perkin Elmer) to dissolve the choroid plexus. For efflux, the choroid plexus was swiftly rinsed in 37°C isotope-free HEPES-aCSF, then transferred into new wells containing 37°C isotope-free HEPES-aCSF with or without 20 μM bumetanide, at 10 s intervals. For each time point, 200 μl of the surrounding HEPES-aCSF was collected into a scintillation vial. At the end of the experiment, the choroid plexus was placed into a scintillation vial containing 200 μl Solvable to dissolve the choroid plexus. Isotope content was determined in 2 ml Ultima Gold^™^ XR scintillation liquid (6013119, Perkin Elmer) using the Tri-Carb 2900TR Liquid Scintillation Analyzer (Packard). The ^86^Rb^+^activity was corrected for extracellular background using ^3^H-mannitol [49, 51]. Efflux data are shown as the natural logarithm of the ^86^Rb^+^ activity at each time point (A_T_) normalized to the initial ^86^Rb^+^ activity (A_0_) as a function of time. The slope from linear regression analysis was used to determine the ^86^Rb^+^ efflux rate constant [49, 51].

### Statistical Analysis

Data analysis and statistical tests were carried out using Graphpad Prism version 9 (GraphPad software). All data were tested for normality of distribution using Shapiro-Wilk test prior to statistical analysis with Student’s unpaired t-test, one way ANOVA with Sidaks multiple comparisons test, and simple linear regression, as stated in figure legends. Data are displayed as the mean and standard error of the mean (SEM) with a p-value < 0.05 employed to define statistical significance. Outliers were determined with Grubbs’ test, when indicated in figure legend.

## RESULTS

### HFD causes elevated ICP in rodents

Obesity in humans is commonly attributed to myriad factors including genetic predisposition, macro- and micro-environmental factors, and lack of physical activity, but the primary recognized feature is consumption of energy rich diet [52]. In order to recapitulate this primary condition in a rodent model, female Wistar rats, aged 6 weeks, were divided into two groups, high fat diet-fed (HFD) and control, and were respectively fed either a HFD (60% Kcal from fat) or a nutrient source-matched control diet (10% Kcal from fat) for 21 weeks. HFD-fed rats gained weight at an accelerated rate (Fig. 1A) and were visibly larger (Fig. 1B) and approximately 18% heavier than their control counterparts on the day of experimentation (HFD: 355 ± 15 g, n = 17 vs control: 301 ± 7 g, n = 14, p < 0.001, Fig 1C), with elevated BMI (HFD: 0.78 ± 0.01 g/cm^2^, n = 17 vs control: 0.66 ± 0.01 g/cm^2^ n = 14, p < 0.001, Fig. 1D). To determine if the HFD rats displayed the elevated ICP characteristic of the IIH patients, the ICP was measured with an epidural pressure probe in the anesthetized rats (Fig. 1E). The ICP was ~ 65% elevated in the HFD-fed rats compared to the control rats (HFD: 5.6 ± 1.2 mmHg, n = 8 vs control: 3.4 ± 1.3 mmHg, n = 8, p < 0.05, Fig. 1F), with the ICP displaying correlation to the body mass across the two groups of experimental rats (n = 16, R^2^ = 0.34, p < 0.05, Fig. 1G). These data demonstrate that dietary-induced body mass elevation can cause the ICP elevation in rodents that is a hallmark feature of IIH in patients. IIH patients display altered pulsatile ICP as evidenced in abnormal ICP mean wave amplitude, possibly arising from IIH-related changes in the neurovasculature of these patients [47]. To determine if such ICP pattern disturbances were present in the HFD-fed rats, ICP waveform analysis quantified the amplitude change of the cardiac waveform (Fig. 1H), revealing elevated mean wave amplitude (MWA) in HFD rats (HFD: 0.18 ± 0.02 mmHg, n = 7 vs control: 0.10 ± 0.02, n = 7 p < 0.05, Fig. 1I). MWA correlates with ICP across the two groups of experimental rats (n = 14, R^2^ = 0.40, p < 0.05, Fig. 1J). We therefore employed this model to determine whether obesity-induced disturbances in CSF dynamics could underlie the elevated ICP in these animals.

**Figure 1.**
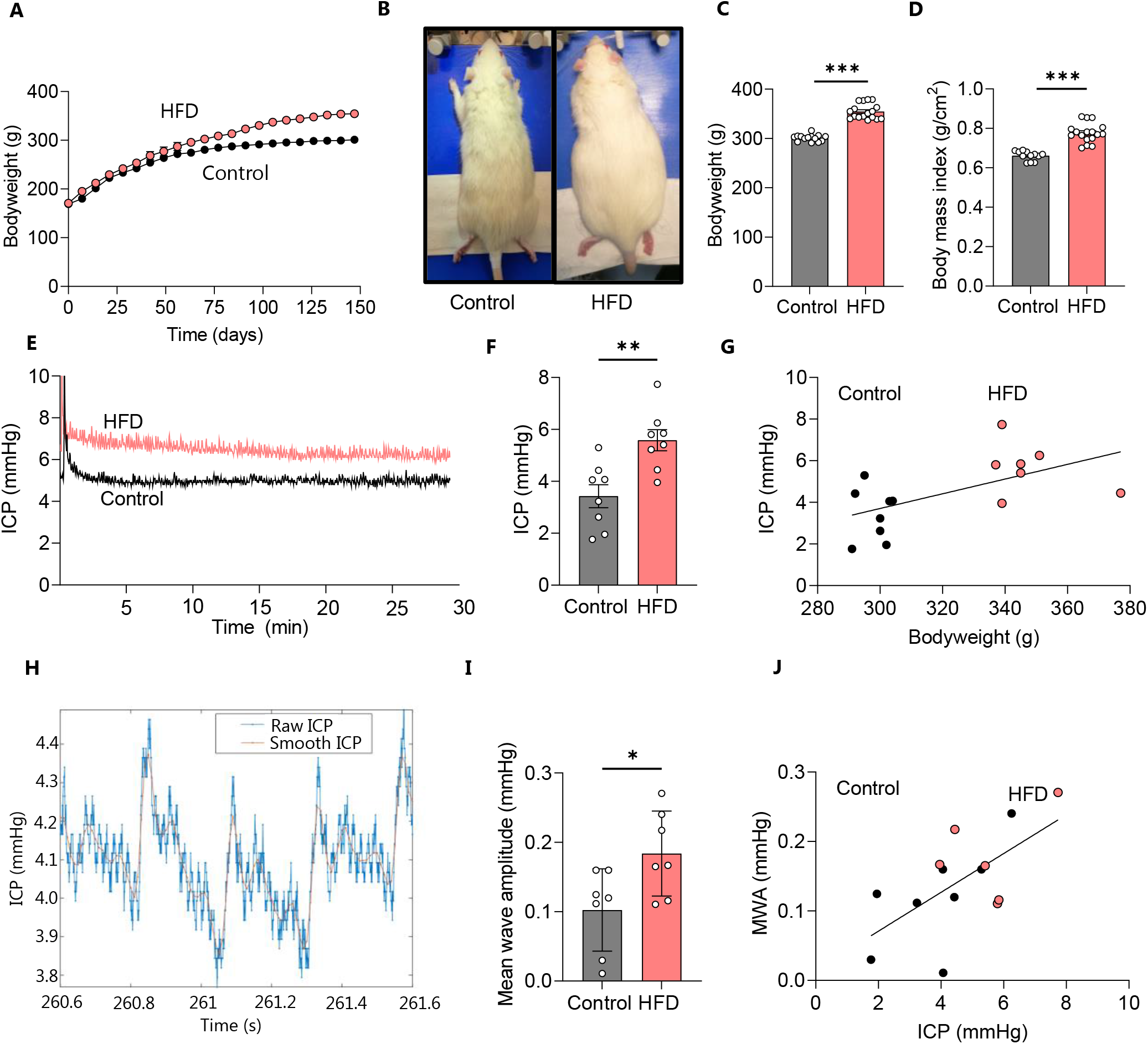
HFD leads to increased bodyweight and intracranial pressure in female rats. **A** Bodyweight increase as a function of time of all control (n = 14) and HFD-fed (n = 17) rats included in the study with the visibly larger rats illustrated in B, the bodyweight at the time of experiments illustrated in **C,** and the BMI at the time of experiment illustrated in **D. E** Representative ICP traces of control and HFD-fed rats with the last 15 min employed for the quantification, summarized in **F. G** ICP correlates with bodyweight across control and HFD-fed rats (R^2^ = 0.34, p < 0.05, n = 16). H Representative ICP trace with “raw” (blue line) and “smoothed” (red line) signals, the latter used to calculate the mean wave amplitude (MWA), represented in I (n = 7 af each). J Correlation analysis of MWA as a function of the ICP, n = 14, R^2^ = 0.40, p < 0.05. Statistical significance evaluated with Student’s unpaired t-test and results shown as mean ± SEM. *p<0.05, **p < 0.01, ***p < 0.001.

### HFD-induced ICP elevation does not arise following elevated CSF production

To determine if HFD-mediated increased ICP arises subsequent to an increased CSF production rate, we assessed this parameter *in vivo* in anesthetized and mechanically ventilated rats with the ventriculo-cisternal perfusion technique [51]. Here, heated and gas-equilibrated aCSF containing a fluorescent dextran is delivered continuously into the lateral ventricle with concomitant fluid collection from a cisterna magna puncture. The dextran dilution taking place during the fluid passage through the ventricles originates from fluid secreted into the ventricles and can thus be employed to calculate the rate of CSF production (Fig. 2A). No significant difference in the rate of CSF production was observed between control and HFD-fed rats (HFD: 5.06 ± 0.91 μl/min, n = 7 vs control: 5.14 ± 1.38 μl/min, n = 6, p = 0.91, Fig. 2B) with no correlation between bodyweight and CSF production rate (n = 13, R^2^ = 0.01, p = 0.73, Fig. 2C). These data indicate that the elevated body weight elicited by HFD is not sufficient to significantly increase CSF production but does not exclude CSF production from a role in modulating ICP. To therefore determine whether the observed increase in ICP associated with excess brain water, the total brain water content was assessed. The total brain weight relative to bodyweight of the HFD rats was significantly lower than control rats (HFD: 5.81 ± 0.16 μg/g, n = 5 vs control: 6.47 ± 0.07 μg/g, n = 5, p < 0.01, Fig. 2D), though this difference is likely attributable to the excess bodyweight of the HFD rats. However, the brain water percentage was not significantly different between the two groups (HFD: 77.6 ± 0.3%, n = 5 vs control: 77.6 ± 0.3%, n = 5, p = 0.16, Fig. 2E). The increase in ICP observed in HFD-fed rats therefore does not appear to arise from an increased brain water content due to elevated CSF secretion rates and may be due to other aspects of CSF dynamics.

**Figure 2.**
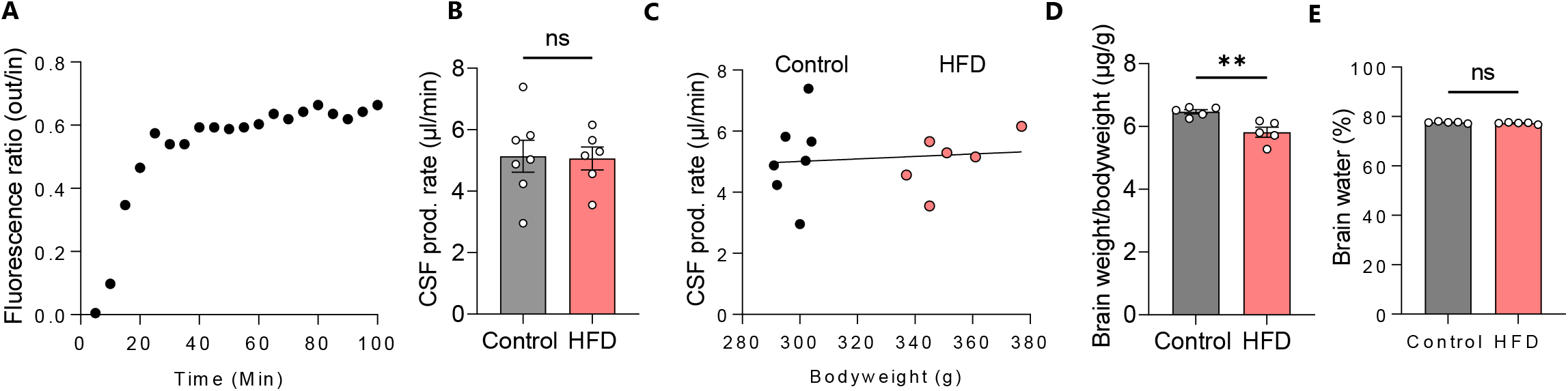
HFD does not cause alterations in CSF production or brain water content. **A** Representative trace of fluorescent dye dilution over the course of a ventriculo-cisternal perfusion assay in a control rat with the final 30 minutes employed for quantification of the CSF production rate illustrated in **B,** n = 6-7. **C** illustrates the CSF production rate as a function of bodyweight, with no significant correlation between the two parameters (n = 6-7, R^2^ = 0.01, p = 0.73). **D** The brain weight relative to the bodyweight of HFD-fed and control rats, n = 5 of each and E the percentage brain water. Statistical significance evaluated with Student’s unpaired t-test and results shown as mean ± SEM. **p<0.01, ns = not significant

### The HFD-mediated ICP elevation is not reflected in overall choroid plexus function

The choroid plexus is the master controller of brain fluid secretion in the mammalian brain [53] and HFD-dependent modulation of its cellular and molecular components could potentially influence brain water dynamics in various ways. To obtain an unbiased manner of revealing potential HFD-mediated changes in functional properties of the choroid plexus, we performed RNA sequencing (RNAseq) of excised choroid plexus obtained from rats fed control diet or HFD. Of the 21,401 expressed genes detected in the choroid plexus (Additional File 1), only 46 of these (0.2%) were differentially expressed between the two groups (Fig. 3A and Additional File 2). A heatmap of the gene expression properties demonstrated that the majority of the differentially expressed genes were downregulated in HFD-fed rats compared to rats receiving control diet (Fig. 3B). A volcano plot demonstrated that of the 46 differentially expressed genes, 33 of these (72%) were downregulated and 13 of these (28%) upregulated (Fig. 3C and Additional File 2). To reveal the functional categories of the differentially expressed genes, we employed GO enrichment analysis (see Methods) to classify 33 of these into protein classes, revealing differentially expressed transcripts categorized as: metabolite interconversion enzymes (39%, 13/33), translational proteins (12%, 4/33), protein-modifying enzymes (9%, 3/33), membrane traffic proteins (9%, 3/33), and transporters (9%, 3/33) – with the remaining 21% scattered over various categories (Fig. 3D). The three differentially expressed transport proteins, ATP5MC1, ATP5MC3, and ATP5PF, though associated with a larger network of differentially expressed genes in the HFD-fed rat choroid plexus (Fig. 3E), are mitochondrial transporters of ATP complex V, not anticipated to be directly associated with CSF formation across the choroid plexus plasma membrane. Overall, the transcript profile of choroid plexus obtained from HFD-fed rats provides no direct indication of a choroid plexipathology as a contributor to the elevated ICP detected in the HFD-fed rats.

**Figure 3.**
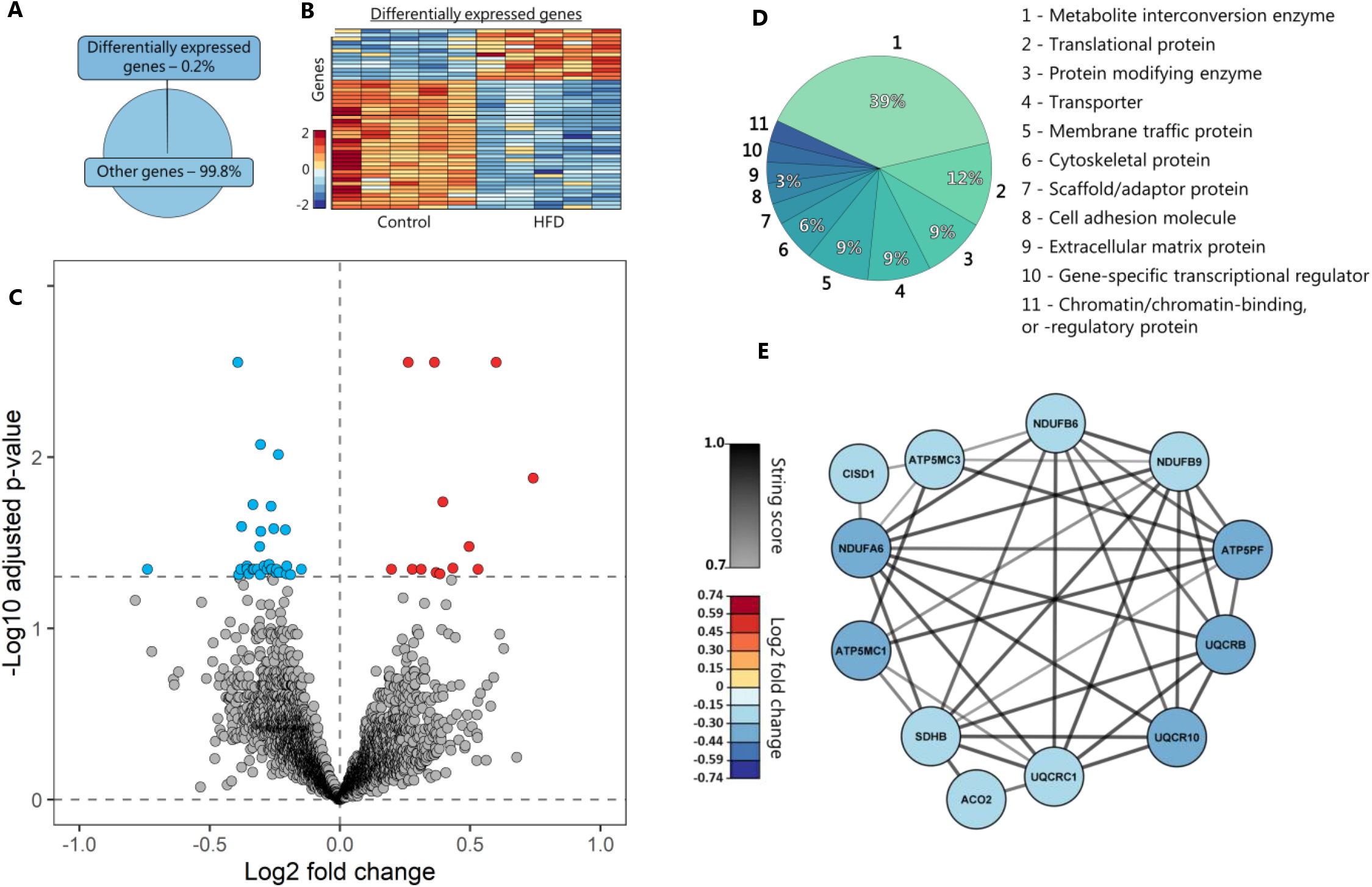
Negligible transcriptomic changes in choroid plexus of HFD-fed rats. **A** illustrates a pie chart of the percentage differentially expressed genes. B A heatmap of the differentially expressed genes in the individual rats, coloured based on the Z-score of normalized fold changes (n = 5 of each). C Volcano plot of all transcripts detected in the choroid plexus of control and HFD-fed rats with differentially expressed genes (Adjusted p-value < 0.05) marked in red (upregulated) or blue (downregulated). **D** GO enrichment analysis of the protein classes. **E** Association protein network analysis of the highest order cluster of genes. The protein nodes are coloured based on log_2_FC (downregulated in blue) and the connecting lines coloured based on the string confidence score from 0.7 (grey) to 1.0 (black).

### CSF androgens as a contributing factor to IIH-related CSF dynamics

To determine whether CSF extracted from the HFD-fed rats differed in testosterone levels compared to their lean counterparts, as observed for IIH patients [33], we performed liquid chromatography-mass spectrometry (LC-MS) analysis on CSF extracted from HFD-fed and control rats. HFD did not induce an elevated testosterone level in the rat CSF (HFD: 1.92 ± 0.03 nmol/l, n = 8 vs control: 1.99 ± 0.04 nmol/l, n = 5, p = 0.17, Fig. 4A), indicating that the HFD-mediated 20% increase in bodyweight alone was not sufficient to cause the hormonal disturbances characteristic of female IIH patients. With the HFD-fed rats not becoming obese, but merely overweight, they may not reach the point at which testosterone dysregulation occurs, and/or such phenomenon may not be linked to bodyweight in the rodent model. Therefore, to determine if elevated CSF testosterone, in itself, could directly modulate CSF dynamics, we subjected lean female Wistar rats to a four-week testosterone treatment regimen [37]. Prior to the initiation of the testosterone regimen, the average weight of the test rats was 142 ± 2 g, n = 51, but the weight of the testosterone-treated (TT) rats increased faster than that of the control rats (Fig. 4B). At the time of experiments, the testosterone-treated rats were visibly larger (Fig. 4C) and significantly heavier than their control counterparts (TT: 136 ± 3.7 g, n = 25 vs control: 92 ± 2.8 g, n = 26, p < 0.001, Fig. 4D). Quantification of plasma and CSF levels of testosterone were performed with LC-MS/MS and revealed elevated testosterone levels in the plasma samples (TT: 100 ± 34 nmol/l, n = 5 vs control: 1.4 ± 0.1 nmol/l, n = 3, p < 0.01, Fig. 4E), but more importantly; also elevated in the CSF (TT: 3.13 ± 1.26 nmol/l, n = 5 vs control: 1.39 ± 0.29 nmol/l, n = 5, p < 0.05, Fig. 4F). The 17β-estradiol level, on the other hand, was not significantly different between the two groups in either the blood (TT: 2.84 ± 0.72 nmol/l, n = 4 vs control: 3.04 ± 0.59 nmol/l, n = 4, p = 0.69, Fig. 4G) or the CSF (TT: 2.70 ± 0.75 nmol/l, n = 4 vs control: 3.04 ± 0.59 nmol/l, n = 5, p = 0.51, Fig. 4H). Of the panel of other steroid hormones and precursors tested during the mass spectrometry analysis, none was significantly elevated in the testosterone-treated rats (Additional File 3).

**Figure 4.**
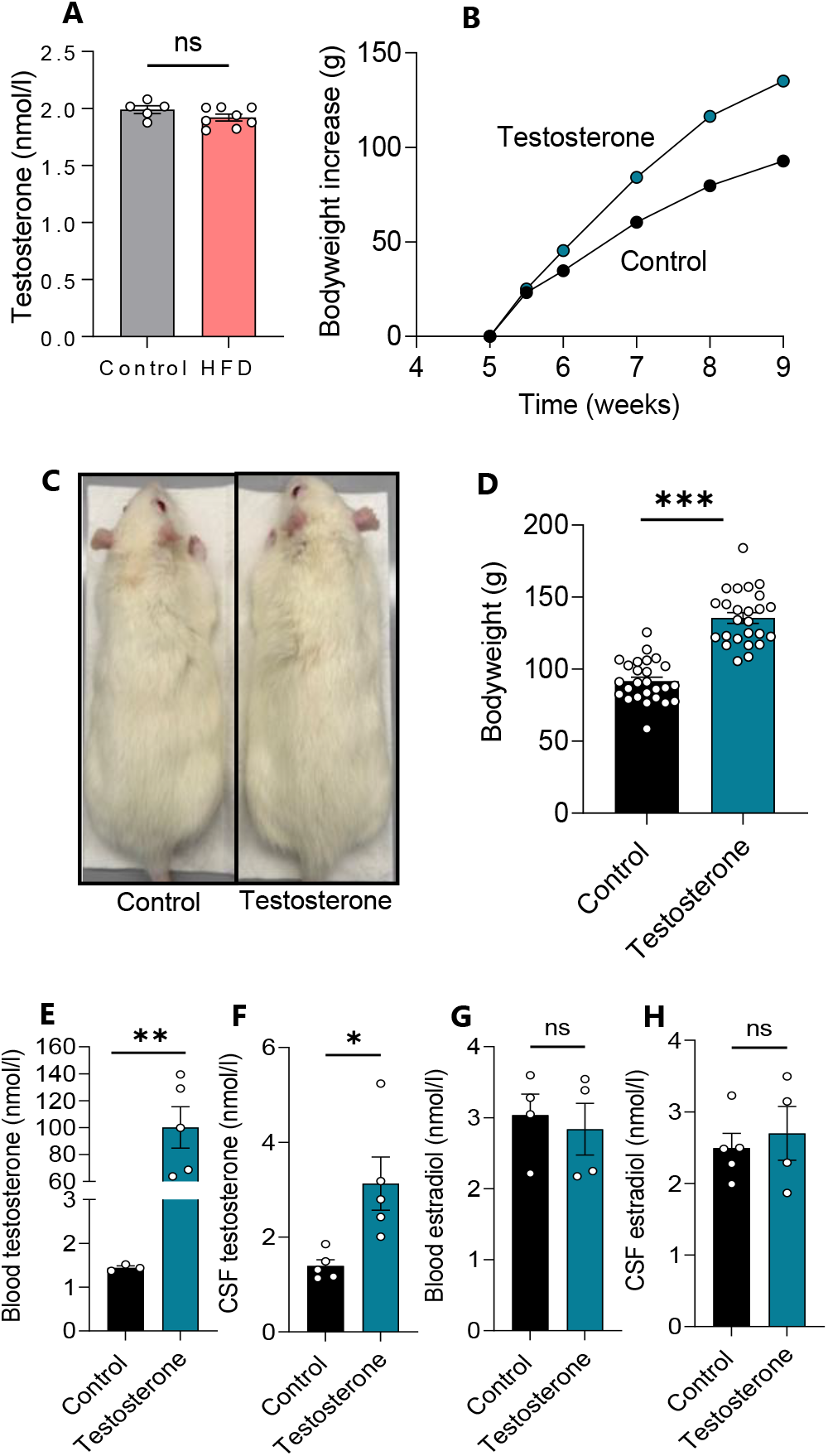
Testosterone treatment increases bodyweight and testosterone levels. **A** CSF levels of testosterone quantified with LC-MS (n = 5-7). **B** Bodyweight increase as a function of time in all rats treated with testosterone (n = 25) or vehicle (n = 26), with C illustrating a visibly larger testosterone-treated rat and **D** their bodyweight at the time of experiments. **E-F** Plasma **(E,** n = 3-5) and CSF **(F,** n = 5) levels of testosterone quantified with LC-MS (after one outlier removed from the control group in E). G-H Plasma (G, n = 4) and CSF (H, n = 4-5) levels of l7β-estradiol quantified with LC-MS (after one outlier removed from the testosterone group in G and H). Statistical significance evaluated with Student’s unpaired t-test and results shown as mean ± SEM. *p < 0.05, **p < 0.01, ***p < 0.001, ns = not significant

To determine if the elevated testosterone modulated the rate of CSF secretion, the LI-COR imaging system was used to acutely assess the CSF flow in testosterone-treated female Wistar rats. This experimental approach relies on injection of a fluorescent dye into the lateral ventricle of the anaesthetized rat, after which the rat is swiftly transferred to the whole animal fluorescent reader and the fluorescence dispersion rate obtained as a proxy for the CSF secretion rate [51, 54–56] (Fig. 5A). The testosterone-treated animals displayed a near doubling of the dye flow (TT: 0.24 ± 0.05 a.u., n = 4 vs control: 0.13 ± 0.05 a.u., n = 5, p < 0.05, Fig. 5B,C) suggesting an elevated rate of CSF secretion. Naïve rats treated acutely with intracerebroventricular testosterone immediately preceding dye injection showed no effects on CSF flow (Additional File 4). To determine whether the testosterone-induced bodyweight increase could contribute to the elevated rate of CSF secretion, we performed correlation analysis of the CSF secretion rate versus the rat bodyweight. With the observed lack of correlation between these two parameters (n = 9, R^2^ = 0.27, p = 0.16, Fig. 5D), the obtained results suggest a testosterone-induced elevation of the CSF secretion rate independently of the rat bodyweight. Such testosterone-induced increase in the rate of CSF secretion could lead to brain water accumulation and thus contribute to the elevated ICP observed in the IIH patients. The brain water content of testosterone-treated rats was assessed with the dry-wet weight technique and revealed no significant difference in the brain water percentages between the two experimental groups (TT: 78.5 ± 0.2%, n = 5 vs control: 78.4 ± 0.3%, n = 5, p = 0.71, Fig. 5E). Although the brain weight was not significantly different in the testosterone-treated rats (TT: 1.89 ± 0.04 g, n = 5 vs control: 7.78 ± 0.03 g, n = 5, p =0.059), the *relative* brain weight was decreased in these animals (TT: 6.81 ± 0.13 μg/g bodyweight, n = 5 vs control: 7.94 ± 0.23 μg/g bodyweight, n = 5, p < 0.01, Fig. 5F), likely due to the excess bodyweight gain induced by testosterone treatment.

**Figure 5.**
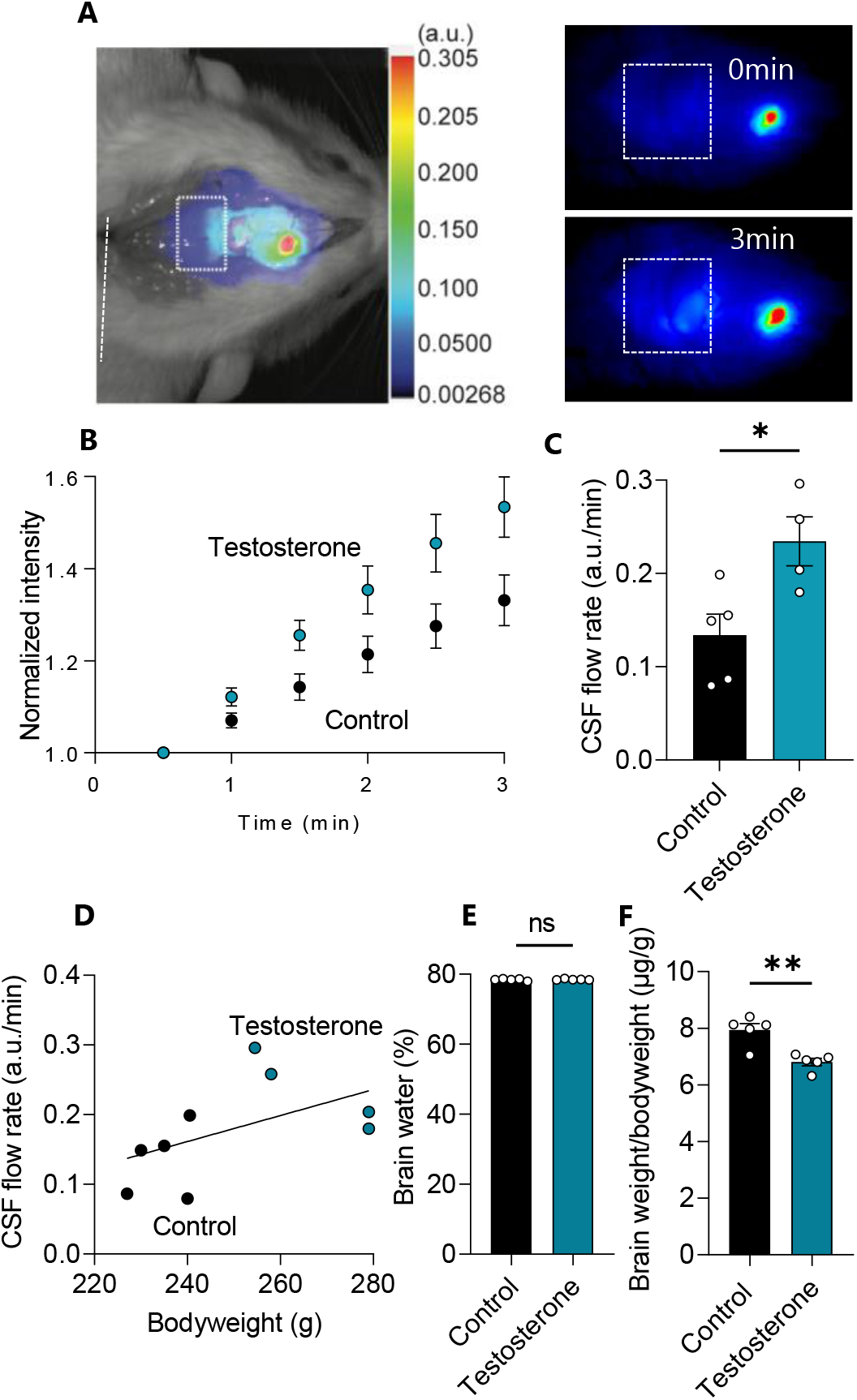
CSF flow increases with testosterone treatment. **A** Representative image of a rat after injection of IRDye 800CW carboxylate dye (superimposed pseudo-color). The square placed in line with lambda indicates the area of dye content quantification, as illustrated in the representative images obtained at t = 0 min and t = 3 min in control rats. B The dye intensity normalized to that obtained in the first image and plotted as a function of time representing flow rate, n = 4-5. **C** Quantification of the dye intensity (flow rate) determined from linear regression in B over the 3 min time window (with one outlier removed from the testosterone group). D The CSF flow rate as a function of bodyweight did not display significant correlation with bodyweight in control and testosterone-treated rats (n = 9, R^2^ = 0.27, p = 0.16). E Percentage brain water in control and testosterone-treated rats (n = 5 of each). F The brain weight relative to the bodyweight of control and testosterone-treated rats (n = 5 of each). Statistical significance evaluated with Student’s unpaired t-test and results shown as mean ± SEM. *p < 0.05, ** p < 0.01, ns = not significant

### Transport activity of NKCC1, not the Na^+^/K^+^-ATPase, is affected by testosterone treatment

To resolve the molecular mechanisms underlying the testosterone-induced elevated CSF secretion, the activity of two choroid plexus transport mechanisms, the Na^+^/K^+^-ATPase and the Na^+^/K^+^/2Cl^-^ cotransporter (NKCC1), both known to contribute to CSF secretion [54] was assessed. To determine the contribution of the Na^+^/K^+^-ATPase to the increased CSF production observed in the testosterone-treated rats, its transport activity was determined in acutely excised choroid plexus from testosterone-treated rats and their control counterparts by influx assays with the radioisotope ^86^Rb^+^, as a congener of K^+^, in the absence and presence of the Na^+^/K^+^-ATPase inhibitor ouabain (Fig. 6A). The ouabain-sensitive fraction of the ^86^Rb^+^ uptake thus represents the Na^+^/K^+^-ATPase activity, which was not significantly different between the two experimental groups (TT: 14.7 ± 6.4 × 10^3^ cpm, n = 4 vs control: 19.2 ± 3.8 × 10^3^ cpm, n = 4, p = 0.25, Fig. 6B). Determination of NKCC1 activity was obtained with an efflux assay of pre-equilibrated ^86^Rb^+^ in the acutely excised choroid plexus in the absence and presence of the NKCC1 inhibitor bumetanide, Fig. 6C-D. The bumetanide-sensitive fraction of the ^86^Rb^+^ efflux thus represents the NKCC1 activity, which was significantly elevated in the choroid plexus obtained from the testosterone-treated rats (TT: 0.38 ± 0.05 min^-1^, n = 6 vs control: 0.32 ± 0.03 min^-1^, n = 5, p < 0.05, Fig. 6E). None of the transport parameters were altered by acute (1 h) exposure of the excised choroid plexus to testosterone (Additional File 4). The data suggest that Na^+^/K^+^-ATPase activation does not underlie the increased CSF flow observed after a four week testosterone treatment in female rats, but that testosterone-induced elevation of NKCC1 transport activity could contribute to the elevated CSF secretion rate observed in these rats.

**Figure 6.**
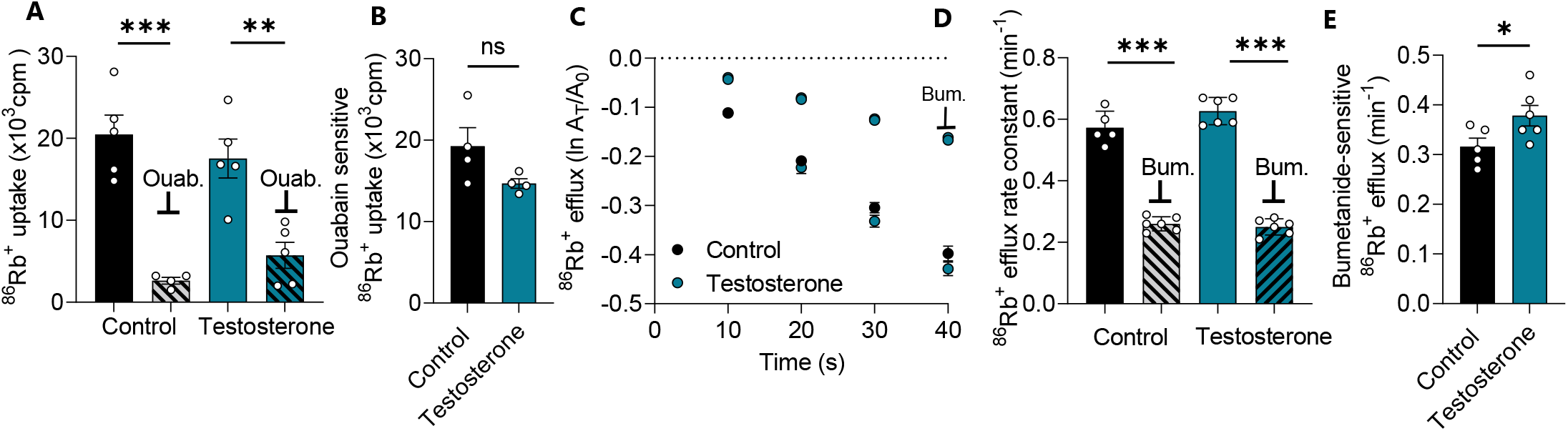
Choroid plexus NKCC1 activity is elevated testosterone-treated rats. **A** ^86^Rb^+^ influx into isolated choroid plexus from control and testosterone-treated rats in the absence and presence of ouabain (Ouab, 2 mM), n = 4-5 (after one outlier removed from the control-ouabain group). B The ouabain-sensitive (Na^+^/K^+^-ATPase-mediated) fraction of the ^86^Rb^+^ influx in choroid plexus from control and testosterone-treated rats (n = 4, after one outlier removed from the testosterone group). C Efflux of ^86^Rb^+^ from choroid plexus obtained from control rats or testosterone-treated rats in the absence or presence of NKCC1 inhibition by 20 μM bumetanide (BUM, n = 5-6 after one outlier removed from the control group). Y-axis is the natural logarithm of the amount left in the choroid plexus at time t (A_t_) divided by the amount at time 0 (A_o_). D The ^86^Rb^+^ efflux rate constant obtained with linear regression of the data from **C. E** The bumetanide-sensitive (NKCC1-mediated) fraction of the ^86^Rb^+^ efflux rate constant obtained from data in D. Results shown as mean ± SEM and statistical significance obtained with Student’s t-test (panels B, E) or ANOVA with Sidaks multiple comparisons test (panel A, D). *p < 0.05,** p < 0.01, *** p < 0.001, ns = not significant

## DISCUSSION

Here, we demonstrate that mimicking of IIH patient characteristics (i.e., female sex, youth, and obesity) in a rat model, led to elevated ICP, the cardinal feature of IIH. The elevated ICP in this rat cohort, however, did not occur with relevant functional changes in the CSF-secreting tissue or with altered CSF secretion rate. However, in rats treated with adjuvant testosterone, causing elevated levels of CSF testosterone (akin to that observed in IIH patients), we observed increased CSF secretion rates. We therefore speculate that CSF testosterone excess is a relevant causal driver in IIH etiology.

Although IIH has been acknowledged as a recognized pathology for more than a century [57, 58], the underlying etiology remains unresolved. According to the Monro-Kellie doctrine, the volume of brain, blood, and CSF is constant. It follows that an increase in any of these compartments, in the absence of a balanced reduction of either of the others, leads to an elevated ICP. Stenosis-induced increased venous blood pressure has thus been proposed as a contributor to the elevated ICP in this patient group [26, 27], although later questioned as stenosis does not seem to correlate with the clinical course of the disease [59, 60]. IIH-related stenosis may thus arise as a consequence of the elevated ICP rather than as its underlying cause [61–63]. The CSF content in IIH patients may [18–20], or may not [64], be elevated and/or redistributed to the subarachnoid space [64] or the brain parenchyma, the latter promoting brain edema and/or slit ventricles [29, 65]. However, subsequent studies failed to detect diffuse brain edema [29, 66] or ventricular slits [64, 65, 67] as obligatory features of IIH patients. The outflow resistance is generally increased in IIH patients [22–24], although not necessarily [68]. With the uncertainty of IIH-related CSF flow disturbances leading to increased ICP and the invasive nature of some of these measurements, we employed a female rat model fed a high fat diet to induce fat gain, as hyperphagia of calorie dense, high fat and palatable food is the principle manner in which human patients become obese [69], to seek to unravel the causative link between overweight and elevated ICP.

HFD-fed rats exhibited increased ICP as a function of their bodyweight, as previously demonstrated (along with other IIH-like symptoms) [35] and aligned with earlier findings in a genetic rat obesity model deficient in the leptin receptor [70], thus manifesting one of the hallmark symptoms of IIH. In addition, the mean wave amplitude of the ICP dynamics was increased in HFD-fed rats and displayed correlation with the ICP elevation observed in these animals. This characteristic mimics the human condition and could indicate that the mean wave amplitude may increase with elevated ICP or, instead, be indicative of microvascular changes in the rat brain tissue similar to those observed in human IIH patients [59, 71]. The observed ICP of the HFD-fed rats was approximately 65% higher than that of the control animals, which aligns with the earlier reports on such rat models [35, 70]. However, this ICP elevation is lower than the (minimum) doubling of the mean lumbar opening pressure observed in IIH patients compared to healthy control subjects [72]. Therefore, although these overweight rats appear to recapitulate the cardinal phenotype of IIH symptomatology, elevated ICP, this model cannot mimic the magnitude of ICP change seen in IIH. This discrepancy may, in part, be due to rodent obesity not being metabolically equivalent to that of human patients [69] and/or to the HFD-fed rats failing to reach the level of obesity (rat BMI roughly translated to a human BMI of 28, 17% increase over their lean counterparts) often observed in IIH patients (average BMI of 32 (range 20-70), 40% increase over their lean counterparts [73]). Similarly, while rat age and developmental cycle does not completely parallel that of humans, the 21 week HFD feeding program of these animals represents a shorter window (roughly equivalent to 12 human years), in which aspects of obesity, such as neuroendocrine imbalances and metabolic inflammation, can exert effects, relative to human IIH patients [74]. At 27 weeks old, rats have a roughly equivalent age of 19 human years, which would represent the early stages of IIH pathophysiology in human patients [74].

Elevated CSF secretion has been proposed to be associated with the elevated ICP in IIH patients [21, 25], although later questioned [68]. The elevated ICP in our HFD-fed rats did not associate with increased brain water content or occur due to an increased rate of CSF secretion, rendering the etiology of the rodent HFD-induced ICP increase unresolved. Our findings contrast those of an earlier report in which rodent CSF secretion appeared increased following a high fat diet (but with no ICP measurements provided) [36]. Although we cannot explain this discrepancy, one possibility could be the degree and rapidity of weight gain, which has been shown to be an exacerbating factor in the presentation of IIH symptoms [73, 75]. In the previous study, female rats increased their weight 3.5-fold over a 7 week course [36], whereas the female rats employed in the current study merely doubled their weight over a 21 week course, an important difference which may affect the appearance of IIH [75]. The small number of experimental animals (n = 3) employed in the previous study, the high volumes of experimentally infused aCSF, and/or the lack of mechanical ventilation of the animals during the prolonged anesthesia required for the procedure [36] may, in addition, contribute to the discrepancy between our findings. In support of the undisturbed CSF secretion observed in the HFD-fed rats, we observed negligible HFD-induced functional alterations in the CSF-secreting tissue choroid plexus, as determined from the transcriptomic profile of this tissue. Merely 46 of the >21,000 transcribed genes obtained from the tissue (0.2%) were differentially expressed in the HFD-fed rat choroid plexus. Of these, only three were categorized as transport proteins, all localized to the mitochondria and all being downregulated, which, taken together, do not support an increase in choroid plexus-mediated CSF secretion in the rats on HFD.

While female sex and obesity are two of the cardinal characteristics of most IIH patients, the great majority of obese females do not have IIH. Striking among biomarkers distinguishing IIH patients from unaffected obese females is elevation of testosterone [33], which is increased to a level over double that of bodyweight-matched controls, but below that of male subjects. The HFD-fed rats did not mimic the elevated CSF testosterone levels, which may be due to their modest bodyweight gain and/or species difference in the response to overweight. Adjuvant testosterone treatment of lean female rats was then employed to mimic the elevated levels of testosterone in the IIH patients. The elevated CSF levels of testosterone caused an increased rate of CSF production in these rats, which was, at least in part, associated with augmented activity of the choroid plexus transport protein, NKCC1, which has been demonstrated as a key contributor to CSF secretion in mice, rats, and dogs [51, 54, 76, 77]. The testosterone-mediated effect on the choroid plexus transport protein and the CSF secretion rate only came about with prolonged treatment with testosterone (4 weeks) and was absent with acute (1 h) testosterone exposure of the excised choroid plexus. In contrast, a previous study with acute testosterone exposure of cultured rodent choroid plexus epithelial cells observed increased Na^+^/K^+^-ATPase activity [33], which we here failed to replicate in the ex *vivo* choroid plexus. Notably, the 17β-estradiol levels were unaffected in both HFD-fed rats and testosterone-treated rats, suggesting that the observed effects on ICP and CSF flow could not be accounted for by l7β-estradiol, as has been proposed as a possible contributor to IIH in patients [32]. However, the rats were not controlled for their oestrous cycle in this study, which may introduce cycle-dependent changes in choroid plexus that may mask testosterone-mediated changes in functionality and is thus considered a limitation to the study. NKCC1 hyperactivity has previously been demonstrated as a regulator of CSF production in another condition of disordered CSF dynamics, i.e., posthemorrhagic hydrocephalus [50, 56, 76], likely occurring with activation of the SPS1-related proline/alanine-rich kinase (SPAK) [56, 76], which is highly expressed in the choroid plexus [78]. The androgen receptor, a nuclear receptor targeted by testosterone [79], is also functionally expressed in the rodent choroid plexus [78, 80, 81]. Androgens induce expression of SPAK [82] and modulate NKCC1 activity [83], which, taken together with the well-established SPAK-dependent modulation of NKCC1 activity [56, 76, 84], provide a potential coupling between the elevated androgenic tone observed in IIH patients and elevated NKCC1-mediated CSF secretion rate. The observed increase in NKCC1 activity in testosterone-treated rats therefore offers a mechanistic explanation for the increased CSF flow observed in these rats. Elevated CSF testosterone in female IIH patients could thus lead to increased CSF production and potentially elevated ICP with prolonged exposure to this androgen, and thereby contribute to the etiology of the disease.

In conclusion, we here demonstrate that although HFD-fed rats gained weight and displayed elevated ICP, their weight gain and ICP increase did not fully reach levels observed in IIH patients. The ICP elevation could not be accounted for by CSF hypersecretion and associated brain fluid accumulation. However, when mimicking the elevated testosterone environment, as observed in patients with IIH, we observed CSF hypersecretion in the experimental rats. Synergic effects of obesity and hyperandrogenism may thus, if sustained for prolonged time and potentially associated with other IIH-promoting factors [28, 59, 72], provide a mechanistic coupling to the elevated ICP observed in the IIH patients. Future *in vivo* animal studies combining obesity and androgen excess may reveal further pieces of the IIH etiology, and androgen blockade, in the context of obesity, may elucidate avenues for therapeutic targeting in IIH.

## Acknowledgements

We are grateful for discussions with Dr RH Jensen, Dr S Eftekhari, and Dr C Westgate regarding the rat feeding regimen, Dr T Toft for assistance with LICOR and brain water experiments, Dr D Barbuskaite for assistance with ICP experiments, and technical assistance from Trine Lind Devantier. We are also grateful for technical assistance from KM Pedersen with mass spectrometry analysis.

## Author contribution

JHW, SNA, ABS, NM designed the research study; JHW, MNJ, SNA carried out the experiments; JHW, SNA, MNJ, JEW, BS analyzed the data; JHW, MNJ, SNA and NM drafted the manuscript and all authors approved the final version.

## Availability of data and materials

The datasets used in the current study are available from the corresponding author on reasonable request. Raw RNAseq data are available at the NCBI GEO database with accession number GSE223582, https://www.ncbi.nlm.nih.gov/geo/query/acc.cgi?acc=GSE223582. Scripts and data analysis are available at: https://github.com/Sorennorge/-MacAulavLab-RNAseq3-Wistar

## Declarations

### Ethics approval

Animal experiments were in compliance with the European Community Council Directive 2010/63/EU on the Protection of Animals used for Scientific Purposes.

### Consent for publication

Not applicable

## Additional Files

**Additional file l.xls** Genes detected in choroid plexus RNAseq from rats fed a control diet and HFD:

**Additional file 2.xls** Differentially expressed genes in choroid plexus RNAseq:

**Additional file 3.pdf** Mass spectrometry analysis of CSF and blood hormones

**Additional file 4.pdf** Acute testosterone treatment does not affect CSF flow or choroid plexus transport rate.

