## Additional file 3&4 for "Modelling idiopathic intracranial hypertension in rats: contributions of high fat diet and testosterone to intracranial pressure and cerebrospinal fluid production"

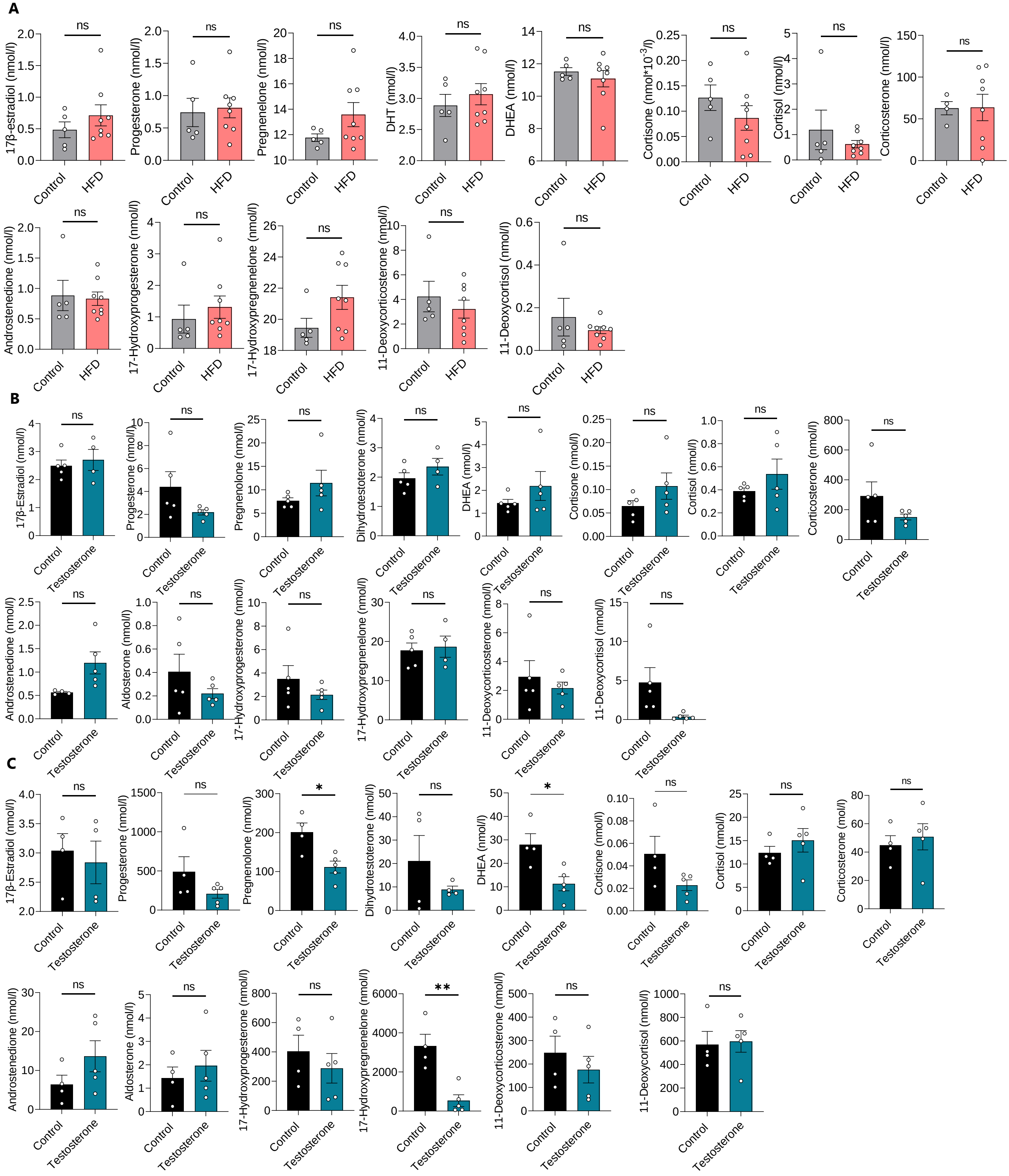


**Additional File 3. Mass spectrometry analysis of CSF and blood hormones: A** CSF hormone levels in control and HFD rats. **B** CSF hormone levels in control and testosterone-treated rats. **C** Blood hormone levels in control and testosterone-treated rats. Pregnenolone and DHEA are significantly decreased in the blood of the testosterone-treated rats, likely due to lack of need for androgen precursors due to downregulation of testosterone production upon treatment with adjuvant testosterone. Results are shown as mean ± SEM. * P <0.05, ** P<0.01, ns = not significant.

### Additional File 4

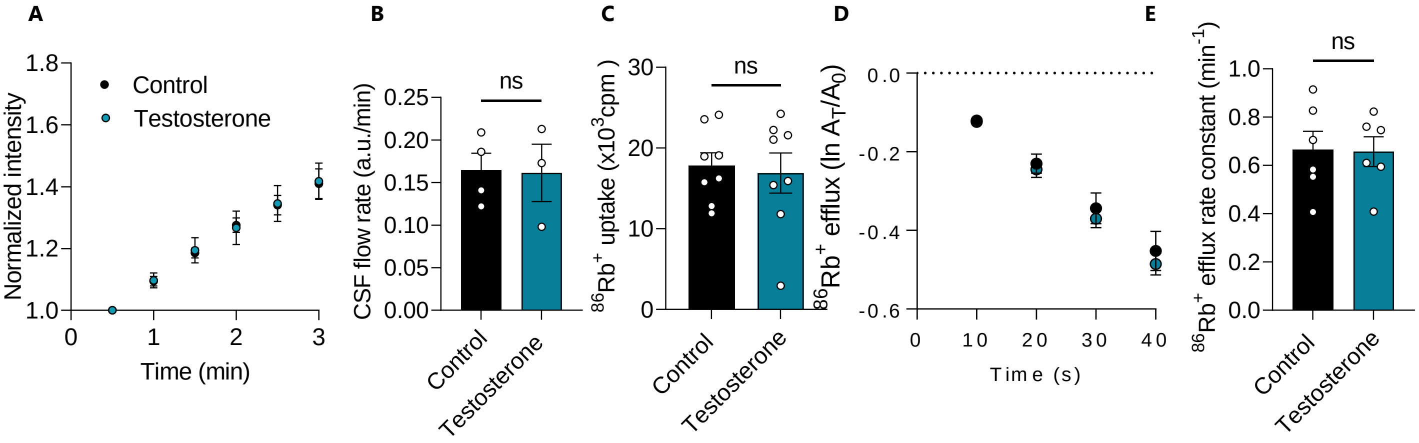


**Additional File 4. Acute testosterone treatment does not affect CSF flow or choroid plexus transport rate. A** The effect of acute intraventricular delivery of testosterone on fluorescent dye flow shown as a function of time. **B** CSF flow rate (expressed as slope of intensity curves in A) is not significantly affected by the acute delivery of testosterone (testosterone: 0.161 ± 0.03 a.u./min, n=3 vs control: 0.165 ± 0.02 a.u./min, n = 4, p=0.94). **C** $^{86}Rb^{+}$ influx in acutely excised choroid plexus after 1 h incubation with 100 nM testosterone shows no significant change compared to control (n = 8). **D+E** $^{86}Rb^{+}$ efflux in acutely excised choroid plexus after 1 h incubation with 100 nM testosterone shows no significant difference compared to control. Results are shown as mean ± SEM. ns = not significant.
